# Behavioral maps organize smartphone interactions in the brain

**DOI:** 10.1101/2025.06.22.660920

**Authors:** Wenyu Wan, K. Richard Ridderinkhof, Arko Ghosh

## Abstract

Smartphone touchscreen interactions are remarkably varied, with each tap and swipe shaped by a distinct combination of cognitive, affective, and environmental factors. One possibility is that the brain plans and processes each interaction anew. Alternatively, the brain may reduce this diversity, processing many distinct interactions through a shared set of recurring neural patterns. The intervals between successive interactions offer one way to detect such shared structure: when interactions are organized by their interval statistics, they reveal regularities that suggest common underlying processes. To test whether these behavioral regularities are mirrored by recurring neural patterns, we recorded EEG during naturalistic smartphone use (n = 53 participants, > 130,000 taps). We placed each interaction on a two-dimensional behavioral map spanning 100 ms to 100 s, defined by its two preceding intervals, and extracted recurring neural patterns time-locked to the touchscreen. Interactions with similar interval context evoked similar neural responses, forming clusters across the behavioral map. Clusters were more pronounced before the interaction than after, suggesting they support upcoming actions. This low-dimensional organization may allow the brain to process diverse interactions through a shared, reusable set of recurring neural patterns rather than planning each anew.

## Introduction

Smartphone use is a hallmark of modern human behavior, with each touchscreen interaction serving a different functional purpose, whether moving to the next picture in an album or typing a message to a friend. Such behavioral diversity poses a computational problem long recognized in motor control: does the brain construct each action anew, or deploy a shared, low-dimensional set of neural processes across diverse actions? Work on muscle synergies and, more recently, on low-dimensional neural population dynamics suggests that varied movements can draw on a common, reused repertoire ^1–3^. A parallel principle holds in perception, where the brain extracts regularities from sensory streams and exploits them to shape ongoing processing ^4–8^. Together these lines suggest a general strategy: capturing high-dimensional structure in lower-dimensional representations that are then deployed to guide behavior. Whether spontaneous, real-world action draws on such a shared, low-dimensional repertoire, as in the varied interactions of everyday smartphone use, remains unknown.

Addressing this research question requires a way to describe the structure of the smartphone use itself. Inspired by Poincaré sections and return maps used to study complex systems, one can treat each touchscreen interaction as a sample from an ongoing behavioral dynamical system rather than as an isolated event ^9–11^. Intricate cascades of neural and cognitive processes underlie behavioral outputs, producing a latent flow: a continuous trajectory through behavioral states that is hard to observe directly. Touchscreen interactions can therefore be viewed as observations of this latent trajectory, analogous to crossings of a Poincaré section ^10^. That is, each interaction is a moment where the flow crosses an observable plane, and each crossing is characterized by the interval since the previous interaction. Following the logic of return maps, which describe how one section-crossing leads to the next, these intervals define a map of next-interval properties, i.e., the joint-interval distribution (JID), a two-dimensional behavioral map of how one inter-interaction interval transitions to the next ^12^. For instance, a short interval followed by another short interval occupies a different location than a short interval followed by a long one. Prior work shows that neighboring behavioral map (JID) locations tend to exhibit coherent behavioral properties: multi-day rhythms, and the effects of neurological disease and aging, each appear in clusters of nearby locations rather than at isolated points ^13–15^. These broad patterns suggest that the brain is not only exposed to interval statistics but may ‘leverage’ them by incorporating these into ongoing processing. This raises the question our study addresses: does neural activity merely reflect these interval statistics as a by-product of behavior, or does it systematically encode this low-dimensional temporal structure? We test this by linking smartphone-derived next-interval statistics to time-resolved EEG.

Prior work offers clues about how brain activity tracks temporal structure, though the intervals studied are typically simpler than those in smartphone behavior and, critically, are imposed as sensory inputs rather than generated by the person’s own actions. Even so, this work establishes that interval tracking is distributed: when monkeys discriminate tactile interval durations, neurons in both somatosensory and prefrontal cortices encode the intervals despite the task engaging a single modality ^16–22^, and brain processes for sub-second and supra-second intervals are partly distinct ^23–26^. We recently asked whether such tracking is organized low-dimensionally, using tactile stimuli with consecutive intervals spanning ∼100 ms to ∼10 s and characterizing the stimulus statistics with the JID framework ^27^. Evoked EEG signals analyzed by next-interval property showed stage-dependent tuning; notably, intermediate stages, putatively in secondary somatosensory cortex, clustered in JID space such that neighboring locations elicited similar responses. This shows that JID-defined interval statistics are associated with structured variation in neural activity albeit only for imposed, sensory intervals. Whether the same holds for intervals arising from spontaneous action remains open.

Here, we investigated the neural correlates of the joint-interval distribution emerging from touchscreen interactions during everyday smartphone use. Time-locked averaging reliably reveals evoked EEG responses around each touch ^28^; we separated these signals according to their JID location, with inter-interaction intervals spanning ∼100 ms to ∼100 s. Using data-driven dimensionality reduction, we extracted prototypical neural patterns distributed across the behavioral map, such that interactions with similar next-interval dynamics were linked to similar signals, forming neuro-behavioral clusters. Analysis was conducted at the individual level: 52 of 53 individuals showed neuro-behavioral clusters in one or more source-separated signals (independent component analysis). Importantly, the neural patterns associated with these clusters peaked at distinct stages around the touchscreen event before dissolving, indicating that interval-specific neural representations are transient, emerging and dissolving at characteristic latencies around the touch.

## Results

### Delineating cortical processes surrounding smartphone interactions based on EEG signal source separation

We first estimated the time window surrounding individual touchscreen interactions that consistently contained neural signals related to the action. To this end, we simultaneously recorded the smartphone interaction-related movement signals with movement sensors attached to the participants’ right thumb and EEG signals (Fig. 1A). The grand-averaged movement signals indicated an onset of kinematic activity ∼ 1 s prior to the smartphone interaction and lasting for another ∼ 1 s after the interaction (Fig.1B, left panel). Although there were considerable inter-individual differences, there was a consistent pattern of downward thumb flexion toward the screen around the time of smartphone touchscreen interaction. In the EEG recordings, the electrode placed over the contralateral somatosensory cortex also showed time-constrained ERP signals (Fig. 1B). To estimate the temporal span of the scalp-wide signals, we used the global field power (GFP), which revealed that the period extending from ∼1.4 s prior to the interaction and ∼ 1 s thereafter (Fig. 1B) contained the strongest brain responses. In keeping with prior observations, these patterns indicate that the discrete smartphone touchscreen interactions are embedded within a temporally defined period of kinematic and neurophysiological activity ^29^. In the subsequent analysis, we focused on this period (Time period of interest, TOI).

**Figure 1.**
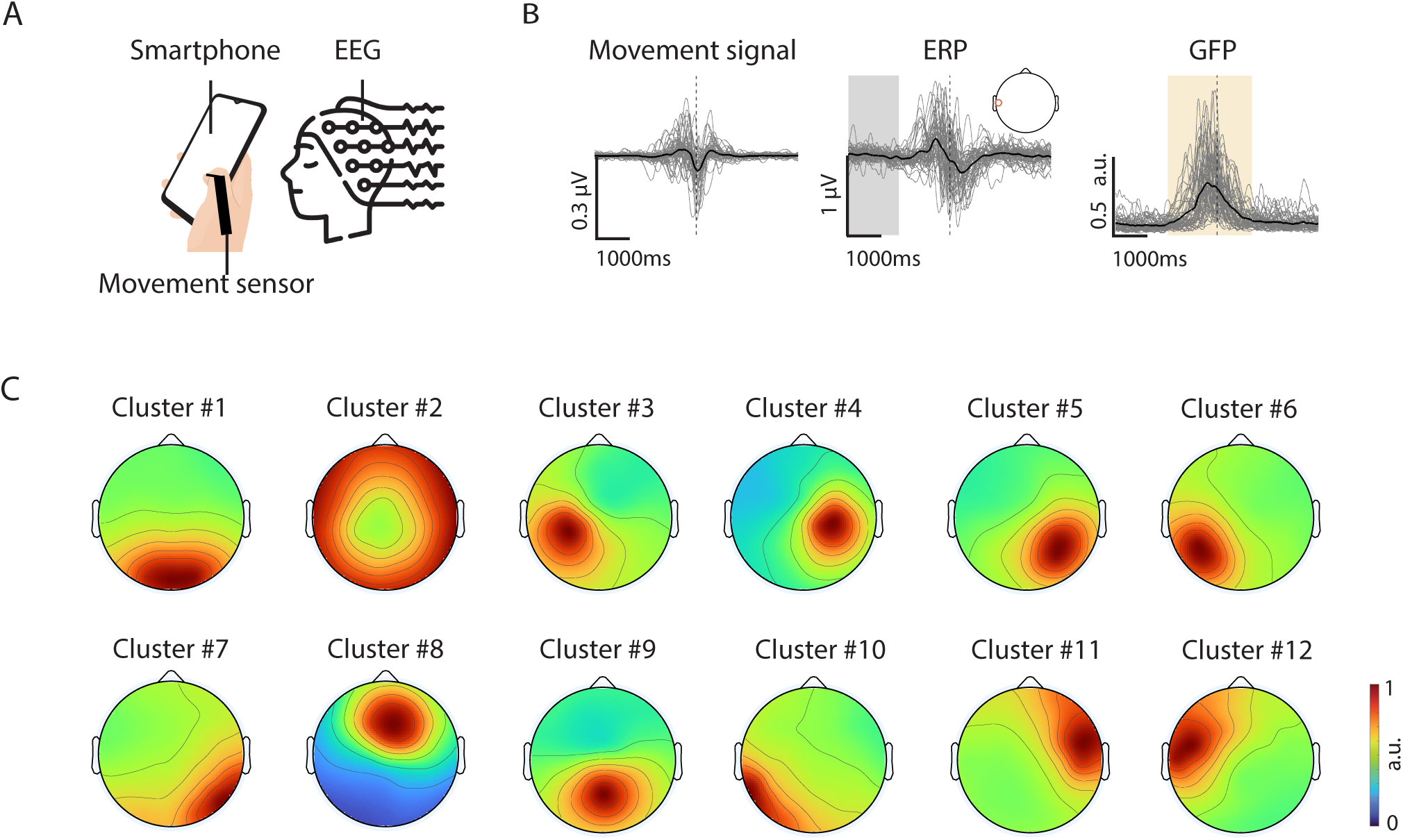
Cortical EEG signals surrounding smartphone touchscreen interactions. (A) Illustration of experimental setup. Timestamps of the smartphone touchscreen event were logged by an App operating in the background on the participant’s personal smartphone. Right thumb flexion-extension was captured by a movement sensor. Cortical population activity was recorded by using 62-channel scalp EEG. (B) Signals time-locked to smartphone touchscreen interactions. Left: black line indicates the grand-averaged (across participants) movement signals (bend sensitive sensor) time locked to the smartphone interaction (horizontal dashed line). Individual movement traces are shown in grey. More negative the value the closer to the smartphone (indicating thumb flexion). Middle: The grand-averaged ERP from an electrode placed over the somatosensory cortex (location shown in the inset) is shown in black while individual ERPs are shown in grey. The analytical baseline period is shaded in grey. Right: black line indicates the grand-averaged global field power (GFP) while grey lines indicate individual’s GFP. The period of the signal used in the subsequent analysis is shaded in yellow (Time period of interest, TOI). (C) Population-wide averaged EEG independent components based on k-means clusters that use the putative dipole locations (n = 433). We identified 12 groups, each representing a putatively distinct brain process and the group order is preserved throughout the manuscript for cross-referencing.

We next separated the putatively distinct brain signal sources captured in the TOI by using independent component analysis (ICA). The analysis at the level of each participant revealed ∼7 sources (median) spanning various cortical locations. To pool information at the population-level, we subsequently grouped these sources (n= 433) according to their putative dipole locations, resulting in 12 typical groups (Fig. 1C). The three most prevalent groups were as follows: group#1 (74 ICs from 47 participants, dipoles centered at MNI coordinates [5,-79,8], located in occipital regions, likely corresponding to visual processing), group#2 (41 ICs from 35 participants, dipoles centered at [-3,-21,22], around the cingulate cortex, likely involved in cognitive and motor control and attentional regulation), group#3 (42 ICs from 31 participants, dipoles centered at [-32,-16,59], around the left central gyrus, likely involved in sensorimotor processing). For the full list of groups, see Supplementary Table 1.

In sum, distinct signal sources were observed surrounding smartphone interactions, implicating processes from motor planning to visual information processing. Our subsequent analysis revealed how these segmented neural signals varied systematically across the behavioral map (JID).

### Cortical signals cluster according to the next-interval smartphone interaction statistics

For each participant, we characterised the next-interval statistics of their smartphone interactions recorded during the EEG measurement session, yielding a two-dimensional behavioral map (JID) per person (JID framework, Fig 2A). Here, the previous smartphone touchscreen interaction interval (say *k*) is represented along with the penultimate interval (say *k*-1). There was substantial variation in these distributions from one person to the next (see Fig 2B). The population average distribution was dominated by next-interval durations of ∼ 1 s. This pattern is in keeping with what is generally observed outside of the laboratory^30^.

**Figure 2.**
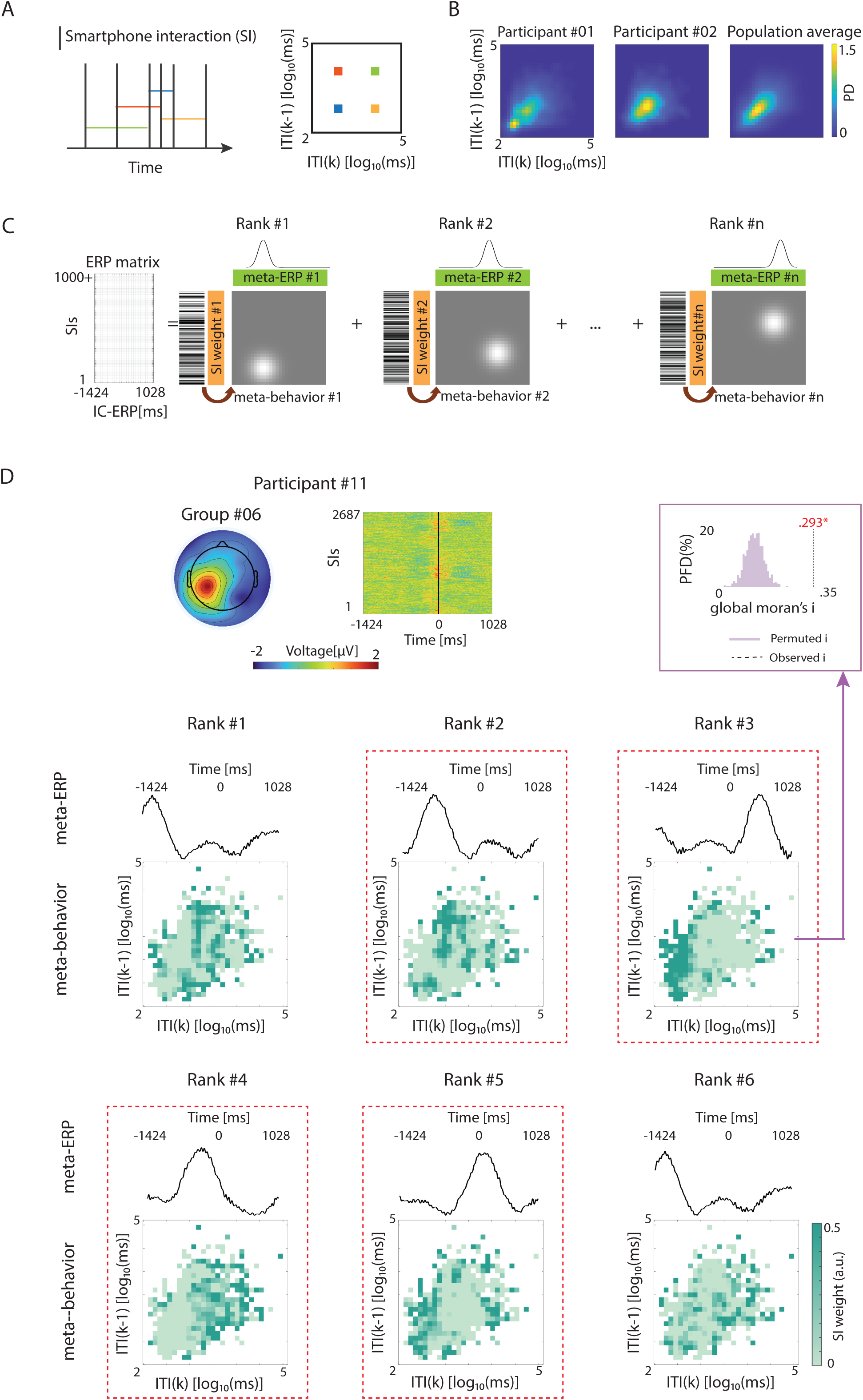
Neuro-behavioral clusters of cortical signals aligned to smartphone next-interval statistics. (A) A schematic explanation of how next-interval statistics of smartphone interactions are captured using a behavioral map: the joint-interval distribution (JID) approach. (B) The next-interval statistics of smartphone interactions were captured during the EEG measurement session. To illustrate the inter-individual differences, the JID from two participants (Sub#01 and #02) are shown, and averaged JID stemming from all the measurement sessions (n = 53). (C) Schematic explanation of parts-based reduction of the neural signals time-locked to smartphone interactions. The source-separated cortical signals were time-locked to the smartphone interactions (SIs) resulting in a signal matrix of SIs x EEG signal latency. We extracted a lower-rank approximation of the matrix using non-negative matrix factorization (NNMF). The reduced prototypical patterns were meta-ERPs – the time-varying neural signal feature – and the corresponding value indicating to what extent the neural feature was present for a given smartphone interaction (meta-SI). The meta-SI values were then projected according to the next-interval statistics of the JID yielding meta-behavior. (D) Example of neuro-behavioral clusters derived from IC Group #06 of Participant #11. The IC-ERP matrix was reduced to 6 (Rank #1 to 6) meta-ERPs and their corresponding meta-behavior (indicating where in JID space each neural pattern was expressed). Each meta-behavior was tested for spatial autocorrelation across the JID using global Moran’s I (an index of whether neighboring locations have similar SI weights). The observed Moran’s I for each meta-behavior was compared to a permutation null, generated by shuffling SI weights across JID locations and recomputing I; meta-behaviour with *p_perm_* <0.05 was considered significantly clustered (boxed in red dashes).

At the individual level, source-separated cortical signals were time-locked to smartphone interactions (SI), yielding a trial-by-trial ERP matrix for each independent component. These matrices exhibited substantial trial-to-trial variation. To identify underlying prototypical patterns, we applied non-negative matrix factorization (NNMF), obtaining a parts-based lower-dimensional representation (Fig. 2C). The parts consisted of prototypical neural response patterns (meta-ERP) and their corresponding SI trial weights (meta-SI). The meta-SI weights were then reprojected onto the JID, yielding a behavioral map of where each pattern was expressed (meta-behavior) and revealing neuro-behavioral clusters.

For instance, in Participant #11 and a source over the sensorimotor cortex (Fig. 2D), a meta-ERP Rank #2, exhibited a neural pattern peaking ∼1 s pre-touch that clustered ∼1 s intervals preceded by ∼10 s long intervals. Another pattern from the same individual and neural source (Rank #3) peaked ∼1 s post touch clustered with intervals <1 s. Global moran’s I confirmed the significant neuro-behavioral clustering (positive values indicate similar values cluster together) relative to permuted null distributions (Fig. 2D). This indicates that specific neural patterns were systematically aligned to behavioral patterns rather than randomly distributed across all behavioral responses.

Across the population, we identified a variety of neuro-behavioral clusters (Fig. 3A), whose exact nature varied across participants and independent components. Most participants (52 out of 53) showed at least one significant neuro-behavioral cluster, with a median of 3 per person (Fig. 3B). Grouping independent components by dipole location, we examined the regional distribution of clustering (Fig. 3B). Regions most frequently showing neuro-behavioral clusters were occipital regions (Group 1, 66% of participants), cingulate cortex (Group 2, 58%), sensorimotor regions (Group 3, 58%), and parietal cortex (Group 9, 67%). Having shown that next-interval statistics are associated with structured variation in neural activity across these distributed regions, we next examined the transient nature of the clusters.

**Figure 3.**
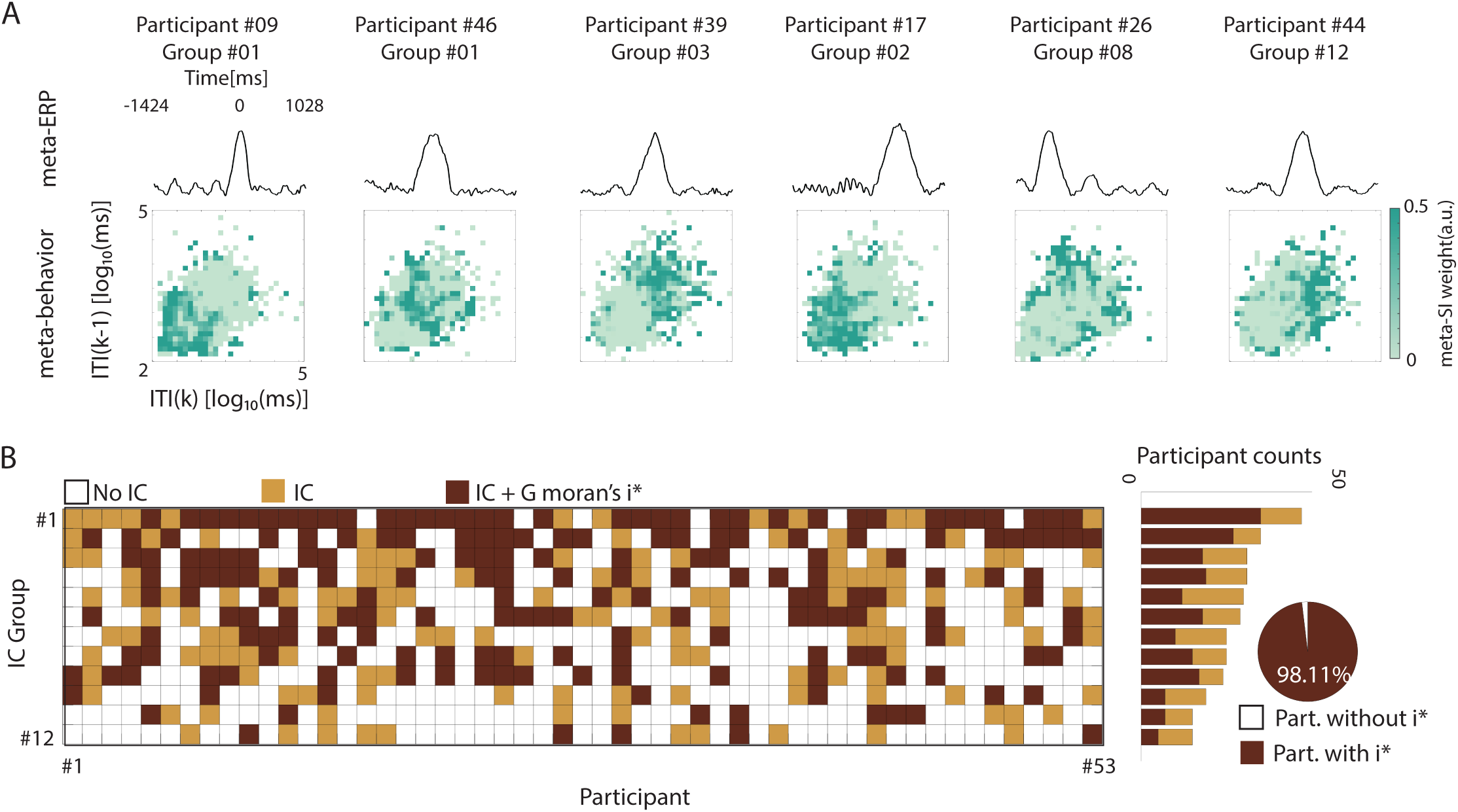
Neuro-behavioral clusters across neural sources and individuals. (A) Across the population we found a range of neuro-behavioral clusters. These example clusters stem from different participants and source-separated cortical processes, showing the NNMF-based lower-dimensional meta-ERPs and their corresponding meta-behaviors. (B) Overview of the source-separated cortical processes found to be clustered in their meta-behaviors (a neural process aligned to next-interval statistics). Each column is a participant and each row a source-separated cortical process (IC group). The brown-filled cell indicates a source-separated cortical signal that showed significant spatial autocorrelation across the behavioral map, and the yellow grid indicates an absence of the same. A white grid indicates the absence of a source-separated process for the corresponding IC group. Bar plots show the number of participants with and without significant clustering for each IC group. The pie chart shows the proportion of participants with significant spatial autocorrelation across the sampled population.

### Timing of the emergent neuro-behavioral clusters relative to smartphone interaction

Our decomposition yields, for each cluster, both its location in behavioral space (meta-behavior) and its temporal profile (meta-ERP). The peak of each meta-ERP marks when that cluster’s neural pattern is most prominent relative to the touch. To characterize this, we analyzed the peak latencies across all putative neural sources and participants.

We calculated the clustering rate for each individual (peak counts/observation duration) and found that the clustering rate was 4.9 clusters per second (SD=3.6) prior to the smartphone interaction and 2.3 (SD=2.1) after the interaction (Fig. 4A). A paired t-test revealed that the clustering rate was significantly higher prior to the interaction than after (*t*(52) = 6.56, *p* = 2.45×10^−8^, 95% CI [1.8 3.4]). The same asymmetry appeared in a complementary analysis that pooled meta-ERPs across participants rather than treating individuals separately (Fig. 4B): the observed proportion of pre-interaction peaks (*p̂*=0.74) exceeded the expected pre-interaction probability (*p*_0_ =0.58) under a uniform temporal null model, as confirmed by a one-sided binomial test (*p*<.05, for the same analysis conducted where the meta-ERPs were grouped according to the putative sources see Supplementary Figure 1). These findings underscore the anticipatory nature of cortical responses to smartphone interactions.

**Figure 4.**
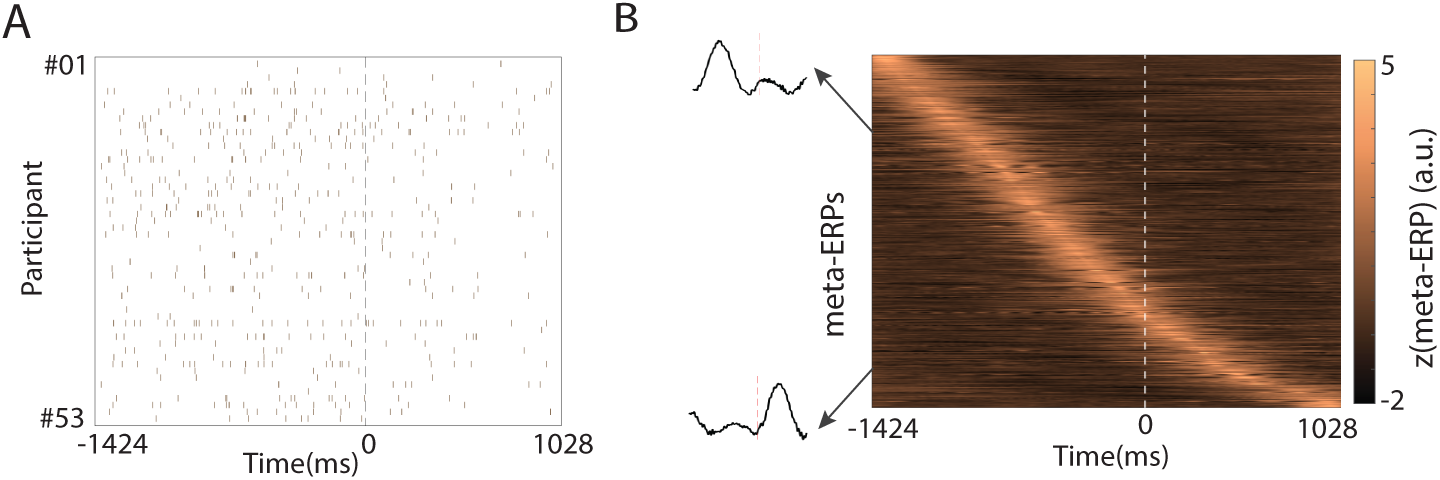
Latency of the neuro-behavioral clusters relative to smartphone interaction. (A) The distribution of meta-ERP peak latencies corresponding to all significant neuro-behavioral clusters observed in each participant (rows). The distribution is time-locked to the smartphone interaction (dashed line). (B) Meta-ERPs pooled from all the participants across all neural processes groups. The data is sorted according to the peak times.

## Discussion

Neural activity surrounding touchscreen interactions is organized into behavioral maps, a pattern observed across participants of a wide age range. Notably, while behaviors with similar interval dynamics (in log-interval space) clustered together, individual clusters were often large, extending across milliseconds to seconds. This suggests that recurring neural patterns can support behaviors spanning multiple temporal scales. These findings suggest a possible mechanism by which neural activity may organize diverse behavioral outputs: interval-based maps form transient, low-dimensional representations that appear predominantly before the interaction, consistent with a role in action preparation.

Our findings are consistent with the value of low-dimensional descriptions in linking brain and behavior, but approach this link from an unconventional direction. A rich literature extracts a low-dimensional representation from neural recordings (for example, neural manifolds derived by dimensionality reduction ^3^) and relates it to behavioral variables such as task onset or type. We establish a complementary approach: the low-dimensional representation is extracted from behavior rather than neural activity, in the form of a behavioral map based on next-interval dynamics, and then linked to neural signals. Because this map is built from spontaneous behavior spanning milliseconds to several seconds, it also extends beyond the sub-second scales typically studied. In designed laboratory tasks with speeded responses, there is a rough separation in neural processing of sub-second and supra-second intervals ^25^. In contrast, some of our clusters spanned multiple timescales, challenging a simple slow-versus-fast separation and suggesting shared neural processes that operate across temporal scales.

The neuro-behavioral clusters were transient, lasting less than a second. This suggests that low-dimensional behavioral maps may play different roles at processing stages surrounding touchscreen interaction. The touchscreen event provided a temporal landmark to examine whether pre-interaction and post-interaction cluster dynamics differed systematically. Prior ERP work suggests that pre-interaction activity is dominated by preparatory sensorimotor processes, whereas post-interaction activity reflects information processing and consolidation ^29^. Together, the transience and the pre/post asymmetry suggest that these low-dimensional maps serve different functions across processing stages: we speculate that preparatory activity uses the recent interval history to guide the upcoming action.

These clusters were observed across the putative signal sources (EEG independent components), including areas associated with motor planning and visual processing. This finding is consistent with the view that temporal structure is encoded across distributed cortical regions ^19,21,25^.

What might underlie the emergence of neurobehavioral clusters? Use-dependent plasticity provides a candidate mechanism: repeated exposure to particular temporal statistics might shape neural organization through experience. Such experience-dependent plasticity could potentially contribute to the substantial inter-individual differences we observed, as each person’s smartphone use generates unique temporal statistics ^14^. This interpretation is consistent with use-dependent cortical plasticity associated with smartphone use ^31,32^, underscores the value of individual-level analysis for real-world neuroscience ^33,34^, and adds to growing evidence for marked individual differences in brain-behavior relationships ^35–39^.

We suggest that transient low-dimensional neural representations may organize behavioural structure according to interval statistics across multiple timescales. A shared set of recurring neural patterns may thus underlie a broad range of behavioral outputs. Future work will need to determine how such temporal maps interact with other neural representations, how they emerge through experience, and what role they play in real-world computation. By capturing behavior as it naturally unfolds, smartphone monitoring offers an unprecedented window into these questions.

## Method

### Participants

Sixty-four participants (age range: 20-81 years; 40 females) were recruited from a participant pool via agestudy.nl research platform ^30^. All participants had no neurological or psychiatric health disorders according to a self-reported screening at enrolment. Four participants were excluded due to a change in self-reported health status: they had developed tinnitus, stroke, arthrosis, or psychiatric disorder. All participants provided written informed consent. This study was approved by the Ethics Committee of Psychology at Leiden University (ERB-reference number: 2020-02-14-Ghosh, dr. A.-V2-2044).

### Smartphone data collection

Smartphone behavior was recorded via the TapCounter app (QuantActions AG, Switzerland) ^40^. All participants accumulated at least one month of smartphone data before EEG measurement. The app logged each interaction timestamp in UTC milliseconds in the background. During EEG recordings, participants freely used one of their four most-used apps from the past month, and switched apps every 10 min after a short break. They were asked to use only their right thumb to touch the screen during measurement. Two experimenters monitored participants using a camera and offered the reminder to use the right thumb when participants deviated. The interactions were stored on a cloud-based data management platform after the EEG measurement. Two participants were excluded because of missing smartphone data on this platform for unknown reasons.

### Movement recording and time alignment

Thumb kinematics was captured using a movement sensor while on the smartphone (Flex Sensor, 112 mm, Digi-Key, Thief River Falls, USA). The sensor was attached to participants’ right thumb’s dorsum using a custom-made jacket. Using this configuration participants were able to freely move their right thumb to interact with smartphone. Analog signals from the movement sensor were sampled at 1 kHz by using a Polybox (Brain Products GmbH, Gilching, Germany) that fed into a digitizer running on the same clock as used for the EEG signals (see below). The continuous movement signal was band-pass filtered between 1 and 10 Hz and then epoched using a 6 s window centered surrounding each interaction and averaged movement signals across trials for each participant.

The smartphone operating system and EEG acquisition system operated on independent clocks requiring post-hoc alignment. We synchronized these clocks using movement kinematics as a bridge signal as described before ^28^. Briefly, movement kinematics (thumb movements) were recorded concurrently with EEG. A pre-trained kinematic-to-touch model predicted touchscreen event times from movement data, successfully identifying the majority of actual events. We then aligned the smartphone and EEG time bases by matching predicted event times (in EEG time) to logged event times (in smartphone time). Four participants were excluded due to poor model performance as confirmed by the average movement sensor signal trace. For participants with multiple recording sessions, the predicted interaction time may differ between the sessions due to subtle kinematic differences impacting the movement profile. We utilized the *alignsignals* function with the *xcorr* method to align the movement waveforms to the waveform whose negative peak occurred closest to the touch time (Matlab’s Signal processing toolbox, MathWorks, Natick, MA, USA). We then calculated the correlation between these time-corrected datasets using Pearson’s correlation coefficient, where high correlation indicates successful alignment across sessions. Datasets with a correlation value below 0.71 were excluded from further analysis. This threshold was determined using the elbow method ^41^ applied to the empirical cumulative distribution (ECDF) of correlation values across participants. This ECDF line showed a clear ‘elbow’ point at r = 0.71, indicating a transition from highly variable to more stable and consistent correlations. Therefore, only datasets with r ≥ 0.71 were retained in further analysis.

### EEG recording and preprocessing

EEG data were collected using a 64-channels active cap (62 scalp electrodes, 2 ocular electrodes) with a customized equidistant layout (ActiCap Snap, EasyCap Gmbh, Wörthsee, Germany). The EEG signals from the active cap system were referenced to the vertex and amplified with a BrainAmp amplifier (Brain Products GmbH, Gilching, Germany), and digitized at a 1 kHz sampling rate. Participants arrived with clean, washed hair and scalp. The scalp contact sites were further degreased using alcohol swabs. Supervisc gel (Easycap GmbH, Herrsching, Germany) was applied to electrically bridge the skin to the electrode, with electrode impedance targeted below 10 kΩ. The data was processed using the EEGLAB toolbox ^42^, signal processing toolbox (MATLAB, Mathworks, Natick) and custom scripts in MATLAB 2023b (MATLAB, Mathworks, Natick). The raw data was high-pass filtered at 0.5 Hz. Inactive smartphone use data segments (pre-interval & post-interval >60 s, for instance resulting from checking in with the participant or offering a drink break) were removed from the analysis.

At the individual level ERPs were computed after low-pass filtering at 20 Hz and removing of the artefacts based on ICA results when over 80% likelihood originating from artifacts like blinks, muscles, etc., as evaluated by ICLabel toolbox ^43^. Cleaned EEG data were then epoched from −3 to 3 s relative to smartphone interaction time. Baseline correction was performed by subtracting the signal mean from −3 to −1.5 s. Grand-averaged ERP signals were then calculated.

To identify the time period of interest (TOI) of event-evoked potentials, we utilized the global field power (GFP), which is a time-variant reference independent indicator of activity based on all of the electrodes ^44^. Here, for each individual, we calculated the averaged ERP for each electrode across trials after trimming extreme value with trimean function. Next, we calculated the standard deviation of ERP across channels at each timepoint (from −3s to 3s with 1ms step). Then, we obtained the grand-averaged GFP (z-transformed) by averaging across participants. To identify the change points of the grand-averaged GFP wave, we utilized the *findchangepts* function with the standard division statistic method and a maximum of two change points (Matlab’s Signal processing toolbox, MathWorks, Natick). The period between these two change points, −1424 and 1048 ms, were defined as the TOI. The IC-ERPs matrix (see below) was limited to this window for subsequent analysis.

### Identifying brain signal sources

We decomposed diverse brain signal sources from the continuous and pre-processed EEG data by adaptive mixture independent component analysis (AMICA) with 2,000 maximum iterations ^45,46^. We estimated the equivalent current dipoles and residual variance for each independent component (IC) using the *DIPFIT* toolbox ^47^. ICs with dipole locations outside of the brain or if the dipole model explained less than 85% of the IC’s scalp voltage distribution (residual variance >15%) were excluded. We further excluded ICs with less than 50% likelihood originating from brain source, as evaluated by the *ICLabel* toolbox ^43^. One participant was excluded due to no surviving IC after screening. Then, the remaining ICs (n=433) from 53 participants were clustered using the *Kmeans* method based on equivalent dipole locations. We explored cluster ranks from 3 to 40, and evaluated the optimal cluster rank using the *evalclusters* function with the ‘Silhouette’ method implemented in MATLAB (Statistics and Machine Learning Toolbox, MathWorks, Natick). Given the variability introduced by random initialization, we repeated this evaluation procedure 100 times and identified the most frequently selected cluster rank as the final optimal cluster rank. To get a reproducible reduction output using the optimal rank approach, we conducted 1,000 repetitions, each with maximum 10,000 iterations. The final reduction results were selected from the repetition with the best sum of squared distance. Source-separated independent component ERP matrix was then reconstructed using *eeg_getdatact (*EEGLAB toolbox, Delorme and Makeig, 2004) based on the cleaned epoched EEG data with the window of TOI, resulting in a rich neural signals matrix of C (components) × T (latencies) × I (smartphone interactions) for each participant.

### Behavioral map: smartphone joint-interval distribution (JID)

Interval durations between smartphone interactions were then calculated, ranging from hundreds of milliseconds to a few seconds. The rich and continuous smartphone behavior can be separated according to the next-interval dynamics using the joint-interval distribution (JID) approach (see Fig. 2A), yielding a behavioral map ^12^. In brief, we established a two-dimensional space where the x-axis represents the pre-interval (say K) while the y-axis represents the penultimate interval (say K-1) in a logarithmic scale spanning from 100 ms to 10 s. We further binned the space into 30 units along the x-axis and y-axis, respectively, resulting in 900 two-dimensional pixels.

### Identifying neuro-behavioral clusters

For any source-separated cortical signal, there was substantial variation from one smartphone interaction to the next, and this was captured in a neural signal matrix organized as the number of interactions x TOI. We extracted the patterns distributed in this rich matrix by using data-driven dimensionality reduction, a variant of non-negative matrix factorization (NNMF, ^48^). The stable and reproducible NNMF (starNNMF) was used to yield time-varying neural features (*meta-ERP*) that occur at different levels across smartphone interactions (*meta-SI*). This method is detailed elsewhere ^49^. In brief, we obtained a non-negative ERP matrix by subtracting the minimum ERP value from the whole matrix. Next, for each participant and source-separated cortical process, we applied the NNMF to the matrix and then searched for the optimal cluster ranks (*c*) between 2 to 15 clusters with 100 repetitions. We identified the optimal *c* which has the minimum error between the real matrix and the reconstructed matrix. Then, with optimal *c* identified, we obtained stable and reproducible decomposition based on 1000 repetitions. Next, the meta-SI was reprojected using the next-interval statistics to reveal how distinct neural processes, activated at specific latencies, map onto the smartphone temporal patterns captured using the JID. For each pixel, we calculated the median value of meta-SIs, thus yielding a meta-behavior representation.

To assess whether the distribution of meta-SI values in the behavioral map (meta-behavior) was randomly patterned or showed two-dimensional autocorrelations, we adopted the global Moran’s I ^50^. This autocorrelation method is typically utilized to identify spatial autocorrelations but the underlying math can be applied to any two-dimensional representation and capture the similarity with its neighborhood ^50^. In brief, we calculated the global Moran’s I for meta-behavior and a corresponding null distribution of I’s based on shuffled ordered trials (1000 permutations). The p-value was estimated as the proportion of permuted Moran’s I values exceeding the observed statistic. The neuro-behavioral clusters with a value of I distinct from the null distribution (*p_perm_*<0.05) was extracted for further analysis.

To address the timing of the neuro-behavioral clusters, we focused on the distribution of meta-ERP peak latencies. We calculated the clustering rate before 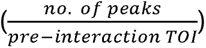 and after 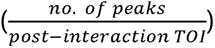 the touchscreen interaction for each individual and contrasted it by using a paired t-test. A separate method was also used to address the same but at the level of each peak instead of each individual. Towards this, we pooled meta-ERPs across participants and estimated the empirical cumulative distribution function (ECDF) of the peak latencies. Here, pre vs. post touch rate differences were assessed using one-sided binomial test (*α* =0.05) to compare the observed proportion of pre-touch peaks 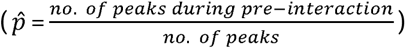 with expected pre-touch probability under a uniform temporal null model 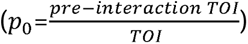. This approach was also used at the level of each IC group i.e., where the meta-ERPs were selectively pooled across the IC groups.

## Data, Materials, and Software Availability

The data will be available on dataverse.nl upon publication. The custom-written scripts used towards this report are shared on: https://github.com/CODELABCODELIB/JID_ERP_Smartphone_2024.

## Acknowledgements

This project was funded by Velux Stiftung (Grant No.1283, awarded to A.G. with Richard Ridderinkhof as co-applicant). We acknowledge Beste Yavuz, Lysanne Groenewegen, Barbora Michalidesová, Lorenzo Van Hoorde for their help with EEG data collection. We also thank all participants for contributing their time and effort. The authors used Claude (Anthropic, Claude Sonnet 4.5 and Opus 4.8, 2026) to improve the clarity, grammar, and organization of the manuscript text. The AI was used solely for language editing; all scientific content, study design, data collection and analysis, interpretation of results, and conclusions were developed by the authors who take full responsibility for the accuracy and integrity of the manuscript.

## Author contributions

A.G. conceived the study. A.G. and W.W. designed the study. W.W acquired, performed EEG pre-processing and analyzed the data with the aid of A.G.. W.W drafted with the aid of A.G. K.R.R., A.G., W.W. edited the manuscript.

---

**Supplementary Figure 1**. The empirical cumulative distribution function (ECDF) of the peak times of the significant neuro-behavioral clusters. The peak times are based on the meta-ERP. (A) Peak times pooled across all participants and neural signal sources. (B-E) The peak times based on specific meta-ERPs pooled from all the participants and the top sources according to the prevalence of neural processes groups (#1, #2, #3, #9; the inserts show scalp locations per cluster).

## Reference

1. Russo, A.A., Bittner, S.R., Perkins, S.M., Seely, J.S., London, B.M., Lara, A.H., Miri, A., Marshall, N.J., Kohn, A., Jessell, T.M., et al. (2018). Motor Cortex Embeds Muscle-like Commands in an Untangled Population Response. Neuron 97, 953–966.e8. 10.1016/j.neuron.2018.01.004.

2. Giszter, S.F. (2015). Motor primitives — new data and future questions. Current Opinion in Neurobiology 33, 156–165. 10.1016/j.conb.2015.04.004.

3. Gallego, J.A., Perich, M.G., Miller, L.E., and Solla, S.A. (2017). Neural Manifolds for the Control of Movement. Neuron 94, 978–984. 10.1016/j.neuron.2017.05.025.

4. Summerfield, C., and de Lange, F.P. (2014). Expectation in perceptual decision making: neural and computational mechanisms. Nat Rev Neurosci 15, 745–756. 10.1038/nrn3838.

5. Saffran, J.R., Aslin, R.N., and Newport, E.L. (1996). Statistical Learning by 8-Month-Old Infants. Science 274, 1926–1928. 10.1126/science.274.5294.1926.

6. Turk-Browne, N.B. (2012). Statistical Learning and Its Consequences. In The Influence of Attention, Learning, and Motivation on Visual Search, M. D. Dodd and J. H. Flowers, eds. (Springer), pp. 117–146. 10.1007/978-1-4614-4794-8_6.

7. Friston, K. (2005). A theory of cortical responses. Philos Trans R Soc Lond B Biol Sci 360, 815–836. 10.1098/rstb.2005.1622.

8. Rao, R.P.N., and Ballard, D.H. (1999). Predictive coding in the visual cortex: a functional interpretation of some extra-classical receptive-field effects. Nat Neurosci 2, 79–87. 10.1038/4580.

9. Fishman, M., Jacono, F.J., Park, S., Jamasebi, R., Thungtong, A., Loparo, K.A., and Dick, T.E. (2012). A method for analyzing temporal patterns of variability of a time series from Poincaré plots. Journal of Applied Physiology 113, 297–306. 10.1152/japplphysiol.01377.2010.

10. Holmes, P. (1990). Poincaré, celestial mechanics, dynamical-systems theory and “chaos.” Physics Reports 193, 137–163. 10.1016/0370-1573(90)90012-Q.

11. Rodieck, R.W., Kiang, N.Y.-S., and Gerstein, G.L. (1962). Some Quantitative Methods for the Study of Spontaneous Activity of Single Neurons. Biophysical Journal 2, 351–368. 10.1016/S0006-3495(62)86860-X.

12. Duckrow, R.B., Ceolini, E., Zaveri, H.P., Brooks, C., and Ghosh, A. (2021). Artificial neural network trained on smartphone behavior can trace epileptiform activity in epilepsy. iScience 24. 10.1016/j.isci.2021.102538.

13. Ceolini, E., Ridderinkhof, K.R., and Ghosh, A. (2024). Age-related behavioral resilience in smartphone touchscreen interaction dynamics. Proceedings of the National Academy of Sciences 121, e2311865121. 10.1073/pnas.2311865121.

14. Ceolini, E., Kock, R., Band, G.P.H., Stoet, G., and Ghosh, A. (2022). Temporal clusters of age-related behavioral alterations captured in smartphone touchscreen interactions. iScience 25, 104791. 10.1016/j.isci.2022.104791.

15. Ceolini, E., and Ghosh, A. (2022). Common multi-day rhythms in smartphone behavior. Preprint at bioRxiv, 10.1101/2022.08.25.505261 https://doi.org/10.1101/2022.08.25.505261.

16. Hernández, A., Nácher, V., Luna, R., Zainos, A., Lemus, L., Alvarez, M., Vázquez, Y., Camarillo, L., and Romo, R. (2010). Decoding a perceptual decision process across cortex. Neuron 66, 300–314.

17. Machens, C.K., Romo, R., and Brody, C.D. (2005). Flexible control of mutual inhibition: a neural model of two-interval discrimination. Science 307, 1121–1124.

18. Bueti, D., Lasaponara, S., Cercignani, M., and Macaluso, E. (2012). Learning about time: plastic changes and interindividual brain differences. Neuron 75, 725–737.

19. Buonomano, D.V., and Laje, R. (2010). Population clocks: motor timing with neural dynamics. Trends in cognitive sciences 14, 520–527.

20. Goel, A., and Buonomano, D.V. (2014). Timing as an intrinsic property of neural networks: evidence from in vivo and in vitro experiments. Philosophical transactions of the Royal Society B: Biological sciences 369, 20120460.

21. Karmarkar, U.R., and Buonomano, D.V. (2007). Timing in the absence of clocks: encoding time in neural network states. Neuron 53, 427–438. 10.1016/j.neuron.2007.01.006.

22. Paton, J.J., and Buonomano, D.V. (2018). The Neural Basis of Timing: Distributed Mechanisms for Diverse Functions. Neuron 98, 687–705. 10.1016/j.neuron.2018.03.045.

23. Koch, G., Oliveri, M., and Caltagirone, C. (2009). Neural networks engaged in milliseconds and seconds time processing: evidence from transcranial magnetic stimulation and patients with cortical or subcortical dysfunction. Philos Trans R Soc Lond B Biol Sci 364, 1907–1918. 10.1098/rstb.2009.0018.

24. Lewis, P.A., and Miall, R.C. (2003). Brain activation patterns during measurement of sub- and supra-second intervals. Neuropsychologia 41, 1583–1592. 10.1016/s0028-3932(03)00118-0.

25. Paton, J.J., and Buonomano, D.V. (2018). The Neural Basis of Timing: Distributed Mechanisms for Diverse Functions. Neuron 98, 687–705. 10.1016/j.neuron.2018.03.045.

26. Petter, E.A., Lusk, N.A., Hesslow, G., and Meck, W.H. (2016). Interactive roles of the cerebellum and striatum in sub-second and supra-second timing: Support for an initiation, continuation, adjustment, and termination (ICAT) model of temporal processing. Neuroscience & Biobehavioral Reviews 71, 739–755.

27. Wan, W., Ridderinkhof, K.R., and Ghosh, A. (2026). The timing of preceding tactile inputs modulates cortical processing. Imaging Neuroscience, in press.

28. Kock, R., Ceolini, E., Groenewegen, L., and Ghosh, A. (2023). Neural processing of goal and non-goal-directed movements on the smartphone. Neuroimage: Reports 3, 100164. 10.1016/j.ynirp.2023.100164.

29. Kock, R., Ceolini, E., Groenewegen, L., and Ghosh, A. (2023). Neural processing of goal and non-goal-directed movements on the smartphone. Neuroimage: Reports 3, 100164. 10.1016/j.ynirp.2023.100164.

30. Ceolini, E., Kock, R., Band, G.P.H., Stoet, G., and Ghosh, A. (2022). Temporal clusters of age-related behavioral alterations captured in smartphone touchscreen interactions. iScience 25. 10.1016/j.isci.2022.104791.

31. Balerna, M., and Ghosh, A. (2018). The details of past actions on a smartphone touchscreen are reflected by intrinsic sensorimotor dynamics. npj Digital Med 1, 1–5. 10.1038/s41746-017-0011-3.

32. Gindrat, A.-D., Chytiris, M., Balerna, M., Rouiller, E.M., and Ghosh, A. (2015). Use-Dependent Cortical Processing from Fingertips in Touchscreen Phone Users. Current Biology 25, 109–116. 10.1016/j.cub.2014.11.026.

33. Akiti, K., Tsutsui-Kimura, I., Xie, Y., Mathis, A., Markowitz, J.E., Anyoha, R., Datta, S.R., Mathis, M.W., Uchida, N., and Watabe-Uchida, M. (2022). Striatal dopamine explains novelty-induced behavioral dynamics and individual variability in threat prediction. Neuron 110, 3789–3804. e9.

34. Levy, D.R., Hunter, N., Lin, S., Robinson, E.M., Gillis, W., Conlin, E.B., Anyoha, R., Shansky, R.M., and Datta, S.R. (2023). Mouse spontaneous behavior reflects individual variation rather than estrous state. Current Biology 33, 1358–1364. e4.

35. Finn, E.S., Shen, X., Scheinost, D., Rosenberg, M.D., Huang, J., Chun, M.M., Papademetris, X., and Constable, R.T. (2015). Functional connectome fingerprinting: identifying individuals using patterns of brain connectivity. Nature neuroscience 18, 1664–1671.

36. Kanai, R., and Rees, G. (2011). The structural basis of inter-individual differences in human behaviour and cognition. Nature Reviews Neuroscience 12, 231–242.

37. Ozdemir, R.A., Tadayon, E., Boucher, P., Momi, D., Karakhanyan, K.A., Fox, M.D., Halko, M.A., Pascual-Leone, A., Shafi, M.M., and Santarnecchi, E. (2020). Individualized perturbation of the human connectome reveals reproducible biomarkers of network dynamics relevant to cognition. Proceedings of the National Academy of Sciences 117, 8115–8125.

38. Wang, J., and He, Y. (2024). Toward individualized connectomes of brain morphology. Trends in Neurosciences 47, 106–119.

39. Zocher, S., Schilling, S., Grzyb, A.N., Adusumilli, V.S., Bogado Lopes, J., Günther, S., Overall, R.W., Winter, Y., and Kempermann, G. (2020). Early-life environmental enrichment generates persistent individualized behavior in mice. Science Advances 6, eabb1478.

40. Balerna, M., and Ghosh, A. (2018). The details of past actions on a smartphone touchscreen are reflected by intrinsic sensorimotor dynamics. npj Digital Med 1, 4. 10.1038/s41746-017-0011-3.

41. Thorndike, R.L. (1953). Who belongs in the family? Psychometrika 18, 267–276.

42. Delorme, A., and Makeig, S. (2004). EEGLAB: an open source toolbox for analysis of single-trial EEG dynamics including independent component analysis. Journal of Neuroscience Methods 134, 9–21. 10.1016/j.jneumeth.2003.10.009.

43. Pion-Tonachini, L., Kreutz-Delgado, K., and Makeig, S. (2019). ICLabel: An automated electroencephalographic independent component classifier, dataset, and website. Neuroimage 198, 181–197.

44. Lehmann, D., and Skrandies, W. (1980). Reference-free identification of components of checkerboard-evoked multichannel potential fields. Electroencephalography and clinical neurophysiology 48, 609–621.

45. Palmer, J.A., Kreutz-Delgado, K., and Makeig, S. (2012). AMICA: An adaptive mixture of independent component analyzers with shared components. Swartz Center for Computatonal Neursoscience, University of California San Diego, Tech. Rep, 1–15.

46. Palmer, J.A., Makeig, S., Kreutz-Delgado, K., and Rao, B.D. (2008). Newton method for the ICA mixture model. In (IEEE), pp. 1805–1808.

47. Delorme, A., Mullen, T., Kothe, C., Acar, Z.A., Bigdely-Shamlo, N., Vankov, A., and Makeig, S. (2011). Research Article EEGLAB, SIFT, NFT, BCILAB, and ERICA: New Tools for Advanced EEG Processing.

48. Lee, D.D., and Seung, H.S. (1999). Learning the parts of objects by non-negative matrix factorization. Nature 401, 788–791. 10.1038/44565.

49. Ceolini, E., and Ghosh, A. (2023). Common multi-day rhythms in smartphone behavior. npj Digit. Med. 6, 49. 10.1038/s41746-023-00799-7.

50. Moran, P.A. (1950). Notes on continuous stochastic phenomena. Biometrika 37, 17–23.

