## Supplementary Table 1 for "Behavioral maps organize smartphone interactions in the brain"

| **cluster** | **n(sub)** | **n(IC)** | **Dipole coordinator**  **(x,y,z)** | **Anatomical label**  **(AAL atlas)** |
| --- | --- | --- | --- | --- |
| #1 | 47 | 74 | 5,-79,8 | Calcarine_R |
| #2 | 35 | 41 | -3,-21,22 | No_label_found |
| #3 | 31 | 42 | -32,-16,59 | Precentral_L |
| #4 | 31 | 45 | 31,-17,-62 | Precentral_R |
| #5 | 30 | 35 | -40,-60,24 | Angular_L |
| #6 | 29 | 36 | 32,-52,38 | Angular_R |
| #7 | 25 | 35 | 42,-52,-19 | Fusiform_R |
| #8 | 25 | 35 | 4,19,46 | Supp_Motor_Area_R |
| #9 | 24 | 32 | -3,-44,61 | Precuneus_L |
| #10 | 19 | 21 | -43,-52,-27 | Cerebellum_Crusl_L |
| #11 | 15 | 20 | 51,0,12 | Rolandic_Oper_R |
| #12 | 15 | 17 | -42,1,5 | Insula_L |

**Supplementary Table. 1.** Summary of 12 source-separated neural processes relative to smartphone interactions.
