## Supplementary figures and images for "Behavioral maps organize smartphone interactions in the brain"

### Supplementary Figure 1

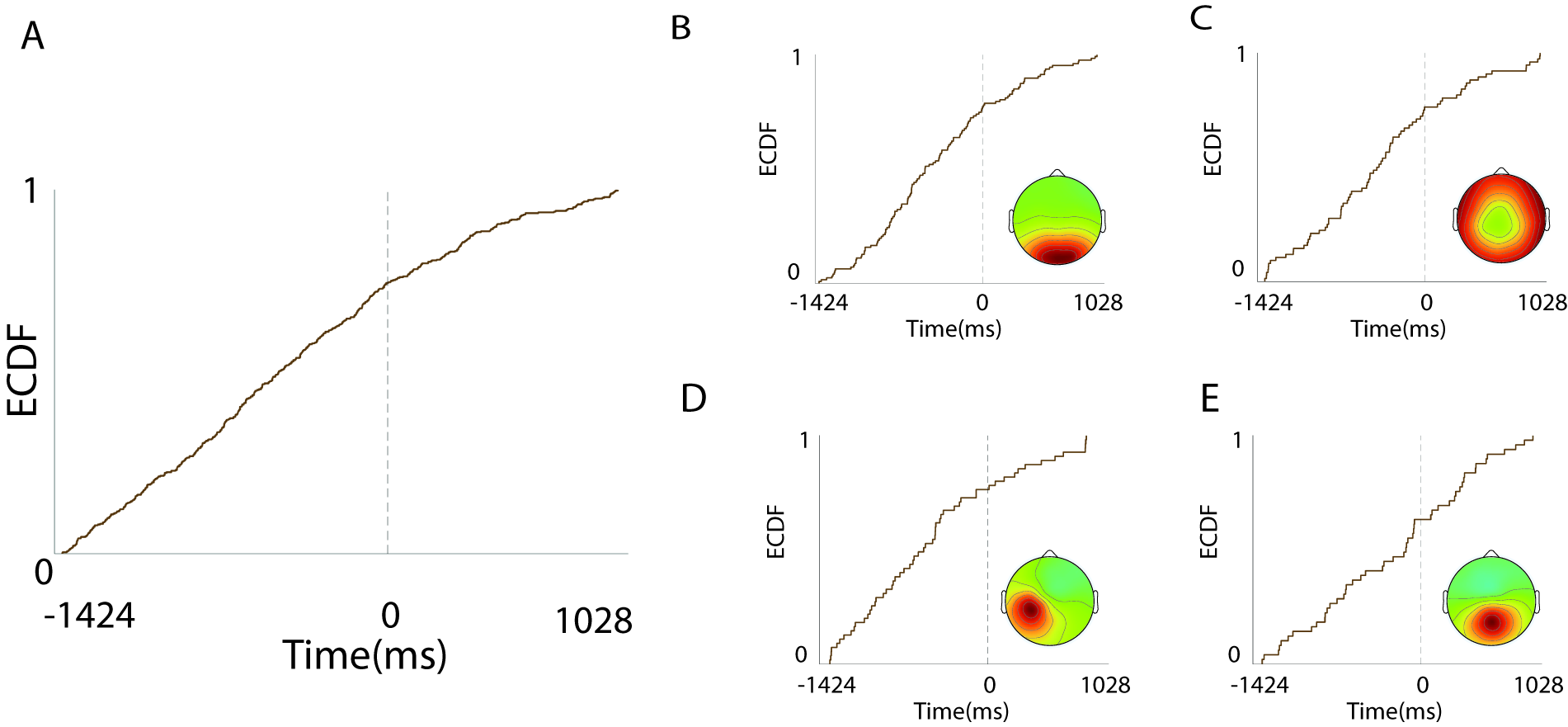
